# Human genetic variation reveals FCRL3 is a lymphocyte receptor for *Yersinia pestis*

**DOI:** 10.1101/2024.12.05.626452

**Authors:** Rachel M. Keener, Sam Shi, Trisha Dalapati, Liuyang Wang, Nicolás M Reinoso-Vizcaino, Micah A. Luftig, Samuel I. Miller, Timothy J. Wilson, Dennis C. Ko

**Author notes:** To whom correspondence should be addressed: Dennis C. Ko, 0049 CARL Building Box 3053, 213 Research Drive, Durham, NC 27710. @denniskoHiHOST.

## Abstract

*Yersinia pestis* is the gram-negative bacterium responsible for plague, one of the deadliest and most feared diseases in human history. This bacterium is known to infect phagocytic cells, such as dendritic cells and macrophages, but interactions with non-phagocytic cells of the adaptive immune system are frequently overlooked despite the importance they likely hold for human infection. To discover human genetic determinants of *Y. pestis* infection, we utilized nearly a thousand genetically diverse lymphoblastoid cell lines in a cellular genome-wide association study method called Hi-HOST (High-throughput Human in-vitrO Susceptibility Testing). We identified a nonsynonymous SNP, rs2282284, in *Fc receptor like 3 (FCRL3*) associated with bacterial invasion of host cells (p=9×10^-8^). *FCRL3* belongs to the immunoglobulin superfamily and is primarily expressed in lymphocytes. rs2282284 is within a tyrosine-based signaling motif, causing an asparagine-to-serine mutation (N721S) in the most common FCRL3 isoform. Overexpression of FCRL3 facilitated attachment and invasion of non-opsonized *Y. pestis.* Additionally, FCRL3 colocalized with *Y. pestis* at sites of cellular attachment, suggesting FCRL3 is a receptor for *Y. pestis.* These properties were variably conserved across the FCRL family, revealing molecular requirements of attachment and invasion, including an Ig-like C2 domain and a SYK interaction motif. Direct binding was confirmed with purified FCRL5 extracellular domain. Following attachment, invasion of *Y. pestis* was dependent on SYK and decreased with the N721S mutation. Unexpectedly, this same variant is associated with risk of chronic hepatitis C virus infection in BioBank Japan. Thus, *Y. pestis* hijacks FCRL proteins, possibly taking advantage of an immune receptor to create a lymphocyte niche during infection.

## Introduction

Pandemics have been potent evolutionary forces throughout human history, selecting for host genetic resistance alleles that likely still impact infectious disease (Fumagalli and Sironi, 2014; Pittman et al., 2016). *Yersinia pestis*, the causative agent of the disease state plague, was responsible for the deadliest pandemic in history – the Black Death of the 14^th^ century (Barbieri et al., 2020). Left untreated, the bubonic plague death rate varied from 30–50% historically (Benedictow, 2004), while the disseminated forms of disease caused almost 100% fatality (Salam et al., 2020). Candidate gene studies (St John et al., 2014; Yang et al., 2019), genome-wide cellular screens (Connor et al., 2018; Klunk et al., 2022; Osei-Owusu et al., 2019) and ancient DNA studies (Immel et al., 2021; Klunk et al., 2022) have implicated various host genes in *Y. pestis* infection. However, a genome-wide association study (GWAS) of *Y. pestis* infection has never been reported.

Genetic association studies have been critical in revealing the genetic architecture of human susceptibility to infectious disease and have also identified host factors that are critical in regulating susceptibility and severity of infection (Gibbs et al., 2022; Hill, 2012). Perhaps the most important host factors in controlling susceptibility to infection are surface receptors that mediate attachment and entry of pathogens. In this regard, human genetic differences affecting entry receptors have been found to confer near complete resistance against two viral infections. Most famously, *CCR5 Δ32* (rs333) is a naturally occurring allele in an HIV-1 co-receptor that protects against HIV-1 infection (Dean et al., 1996; Liu et al., 1996; Samson et al., 1996). Indeed, the first three individuals who have been cured of HIV-1 infection received stem cell transplants from homozygous *CCR5 Δ32* donors (Hutter et al., 2009; Jensen et al., 2023). Similarly, a nonsense mutation in *FUT2* (rs601338) results in the absence of specific fucosylated oligosaccharides and a loss of susceptibility to infection with certain Norwalk virus strains that use the sugars to attach to host cells (Lindesmith et al., 2003). Thus, there is strong precedent for human genetic variation impacting viral entry receptors to confer human resistance, but to our knowledge this has not been reported for bacterial pathogens.

In *Yersinia* species that infect humans, there are several cell-type specific host receptors that contribute to disease. However, while the characterization of binding of *Y. pseudotuberculosis* invasin to host beta-integrin were landmark studies in understanding bacterial entry into host cells (Isberg and Leong, 1990; Isberg et al., 1987), invasin is a pseudogene in *Y. pestis* (Parkhill et al., 2001). Similarly, the YadA adhesin is also a pseudogene in *Y. pestis* (Parkhill et al., 2001). Most notably, FPR1 was shown to contribute to species-specific disease as a receptor for the *Y. pestis* type 3 secretion system (T3SS) (Osei-Owusu et al., 2019), and CD205 was reported to serve as a dendritic cell entry receptor (Zhang et al., 2008). Dendritic cells have been shown to traffic *Y.* pestis to the draining lymph node and help spread bacteria through the lymphatic system (Arifuzzaman et al., 2018; St John et al., 2014). Most work on *Y. pestis*-host interactions have focused on neutrophils, macrophages, and dendritic cells without more closely examining a large component of lymphatic organs: lymphoid lineage cells. While *Y. pestis* is found to alter gene expression and interact with these cells during infection (Klunk et al., 2022; Li and Yang, 2008; Zhao et al., 2023), there are no recognized *Y. pestis* receptors specific to B and T cells.

Here, we used a high-throughput cellular GWAS approach called Hi-HOST (High-Throughput Human In-vitrO Susceptibility Testing) (Gibbs et al., 2023; Ko et al., 2009; Wang et al., 2018a) to model a “pandemic-in-a-plate” and harness global natural human genetic diversity to discover variation that regulates *Y. pestis* invasion into lymphoblastoid cell lines (LCLs; EBV-immortalized B cells). We identified a nonsynonymous variant (rs2282284) in *Fc receptor like 3* (*FCRL3*) associated with cell entry of unopsonized *Y. pestis*. We show that FCRL3 and other FCRL proteins facilitate attachment and invasion of *Y. pestis* and that rs2282284 severely impairs a motif that influences SYK-mediated invasion of human cells. Thus, human genetic variation has revealed a lymphocyte-specific receptor for *Y. pestis*.

## Results

### A cellular GWAS of *Yersinia pestis* infection

To uncover human genetic differences that impact interactions with *Y. pestis*, we applied Hi-HOST to *Y. pestis* host cell invasion and early survival (**Fig. 1A).** This system presented an opportunity to identify common (>1%) human variants that impact general host factors in *Y. pestis* infection, as well as specific factors that may play a critical role in the lymphatic tropism of disease. Briefly, a modified gentamicin-protection assay using *Y. pestis* KIM6+ tagged with isopropyl ß-D-1-thiogalactopyranoside (IPTG)-inducible GFP was coupled with flow-cytometric quantification of the percentage of GFP+ cells at 4 hrs post-infection (hpi; referred to as “invasion”). KIM6+ lacks the pYV/pCD1 virulence plasmid that encodes the T3SS that is known to reduce uptake into host cells (Burrows and Bacon, 1956; Fallman et al., 2002; Janssen et al., 1963; Spinner et al., 2008), creating an environment similar to early mammalian infection before activation of the *Yersinia* virulence plasmid. Notably, invasion of *Y. pestis* was severely inhibited by the presence of serum (**Fig. S1A**), so the assay was conducted in serum-free media. Hi-HOST was carried out on 961 HapMap (Consortium, 2005; Frazer et al., 2007) and 1000 Genomes Project (Genomes Project et al., 2012; Genomes Project et al., 2015) LCLs from 8 global populations and each LCL was measured in three independent experiments (**Table S1**). Inter-individual variation was highly reproducible (repeatability = 0.90, CI = [0.89, 0.91]) and a substantial fraction of this variation could be attributed to genetic differences based on two methods of heritability estimation (h^2^ = 0.15 by parent-offspring regression; h^2^ = 0.19 by genome-wide SNPs; **Fig. S1B**).

**Figure 1.**
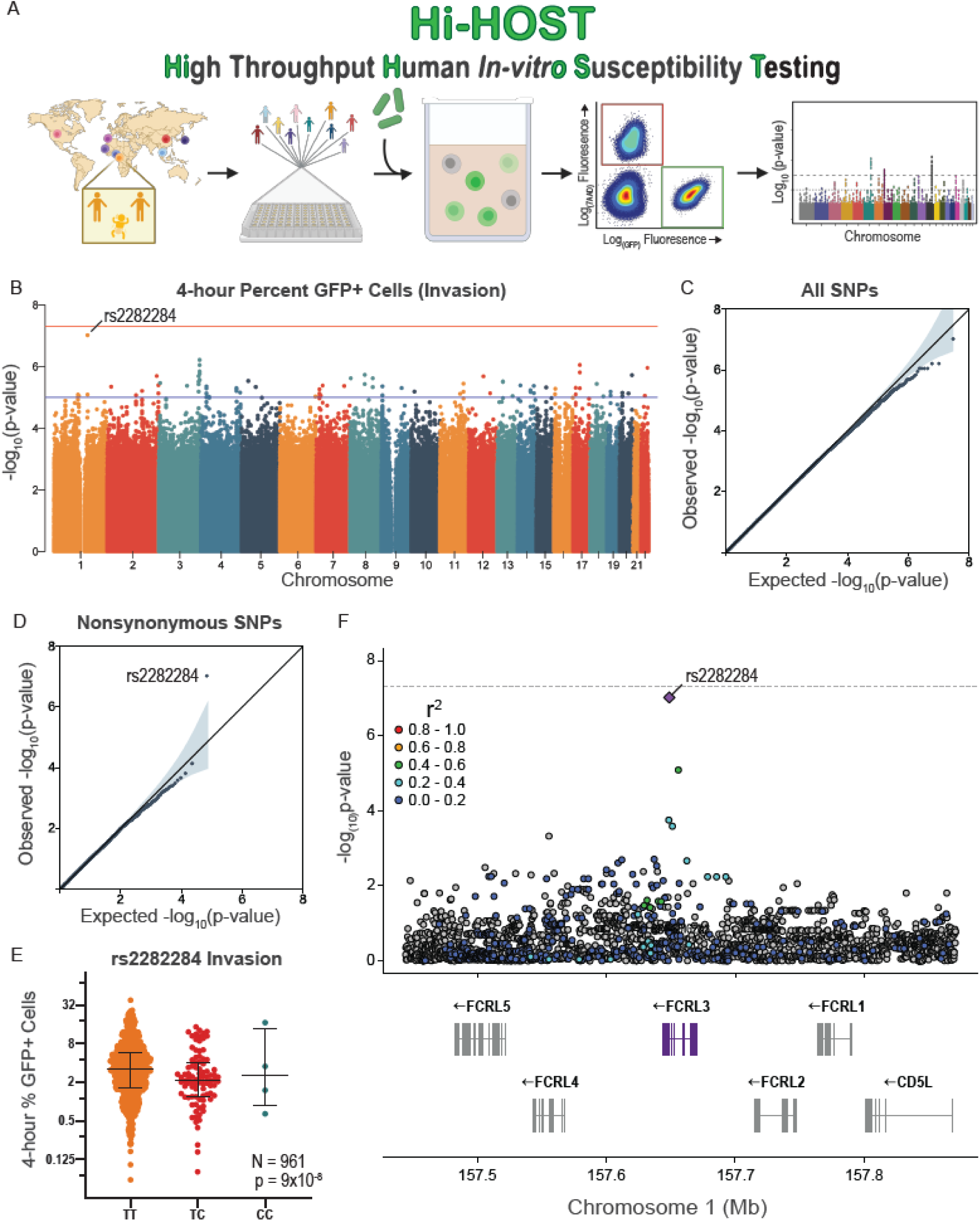
Hi-HOST cellular GWAS of *Y. pestis* invasion reveals the genetic architecture of human host cell interactions with the causative agent of bubonic plague. (**A**) Flow chart of Hi-HOST for *Y. pestis* invasion. The map displays the 8 populations tested in this study (CHB, KHV, JPT, GWD, YRI, ESN, CEU, IBS) (**B**) Manhattan plot of *Y. pestis* invasion measured at 4 hpi for 961 genetically diverse LCLs after analysis with QFAM-parents using PLINK. Orange and blue lines show suggestive (p=1×10^-5^) and genome-wide significant (p=5×10^-8^) thresholds. The lead SNP on chromosome 1 is rs2282284. (**C**) Quantile-quantile plot of observed vs. expected –log(p-values) for all 15,213,612 SNPs demonstrates minimal deviation from the null distribution (grey line). (**D**) Stratified quantile-quantile plot for 33,723 nonsynonymous SNPs reveals a single SNP with a p-value lower than expected by chance (rs2282284; p=9×10^-8^). (**E**) Genotypic mean plot of rs2282284 for *Y. pestis* invasion in 961 LCLs. Bars indicate the mean and interquartile range. The C allele is associated with a lower percentage of infected cells at 4 hrs at p = 9×10^-8^ by QFAM-parents analysis. (**F**) A local Manhattan plot displaying a 500 kb region of chromosome 1 demonstrates that rs2282284 is within the *FCRL3* gene on chromosome 1. A purple diamond denotes rs2282284 and LD with SNPs in the locus is shown by red >= 0.8, orange = 0.6-0.8, green = 0.4-0.6, light blue = 0.2-0.4, dark blue < 0.2, and grey has no LD data. LD was determined by all populations in the study based on combined data from LDLink accessed through “*locuszoomr*” R package.

We conducted a family-based GWAS on all variants with minor allele frequency (MAF) > 1% (QFAM-parents in PLINK; (Purcell et al., 2007; Purcell et al., 2005), leveraging parent-offspring trios to control for population structure. Analysis of the quantile-quantile plot revealed no deviation of –log_10_(p-values) from the null distribution (λ = 1). While there was an absence of genome-wide significant hits (p < 5 x 10^-8^; **Fig. 1B, C**), stratifying GWAS by variant annotations that are more likely to be functional (e.g. nonsynonymous, 5’ or 3’ UTR, expression quantitative trait loci (eQTL)) has proven to be an effective strategy for finding true positive associations (Schork et al., 2013). In fact, stratifying for nonsynonymous variants revealed a single SNP, rs2282284, highly deviated from the expected null distribution (**Fig. 1D**). Here, the minor allele C was associated with lower invasion of *Y. pestis* (p=9×10^-8^, **Fig. 1E**). Focusing on the locus, this nonsynonymous SNP is within *Fc receptor like 3* (*FCRL3*), a gene encoding a member of the FCRL family (Davis, 2007; Tolnay, 2022) in a region on chromosome 1 that has undergone paralogous expansion (**Fig. 1F**). Stratification of the GWAS by continental ancestry showed the phenotype at rs2282284 was primarily driven by the African-ancestry LCLs (due to the higher frequency of the minor C allele), although the directionality of effect was the same in all populations (**Fig. S2**). Interestingly, *FCRL3* is moderately induced during *Y. pestis* infection of primary blood mononuclear cells, specifically B cells (1.19-fold at 5 hrs, p=0.0019; (Klunk et al., 2022)).

### FCRL3 stimulates *Y. pestis* attachment and invasion

FCRLs are a family of type I transmembrane glycoproteins expressed primarily on B cells (Davis, 2007; Tolnay, 2022). While they are structurally similar to the antibody binding Fc receptors (FcRs), only recently has their direct binding capacity to specific Ig classes been biochemically demonstrated (Agarwal et al., 2020; Wilson et al., 2012). Other FcRs have been reported to serve as entry receptors for cytomegalovirus and Enterovirus B (Maidji et al., 2006; Zhao et al., 2019) and phagocytosis of opsonized bacteria can also be triggered by FcRs (Charles A Janeway, 2001; Indik et al., 1995a; Laassili et al., 2023). However, invasion of LCLs by *Y. pestis* in Hi-HOST specifically occurs in the absence of serum, meaning the bacteria is unopsonized during interaction with FCRL3. Following ligand binding, Fc receptors utilize immunoreceptor tyrosine-based signaling motifs to trigger cellular processes, including receptor-mediated endocytosis and phagocytosis (Getahun and Cambier, 2015; Indik et al., 1995b; Tay et al., 2019; Xu et al., 2002). FCRL3 has one immunoreceptor tyrosine-based activation motif (ITAM; amino acids 650-665: Y_650_SNVNPGDSNPIY_662_SQI) that is known to activate internalization in other Fc receptors (Crowley et al., 1997), and one immunoreceptor tyrosine-based inhibitory motif (ITIM; amino acids 690-695: VLYSEL) (**Fig. 2A**) (Xu et al., 2002). Additionally, the nonsynonymous SNP rs2282284 causes an asparagine to serine change at FCRL3 residue 721, directly next to an additional phospho-tyrosine (Y722) in an ITIM-like motif (amino acids 719-725: EENYENV), which is thought to associate with Src homology 2 domain-containing phosphatases but does not completely recapitulate ITIM phenotypes or sequence homology (Kochi et al., 2009; Xu et al., 2002). In fact, this motif may rather be acting as an ITAM-like motif, perhaps in the way of a HemITAM (DED/EGYxxL) which are highly plastic in their function based on the amino acids they contain (Bauer and Steinle, 2017).Therefore, we hypothesized that FCRL3 might be serving as a cell-surface receptor for *Y. pestis,* and rs2282284 might interfere with cytosolic signaling required to mediate uptake of FCRL3 bound to *Y. pestis*.

**Figure 2.**
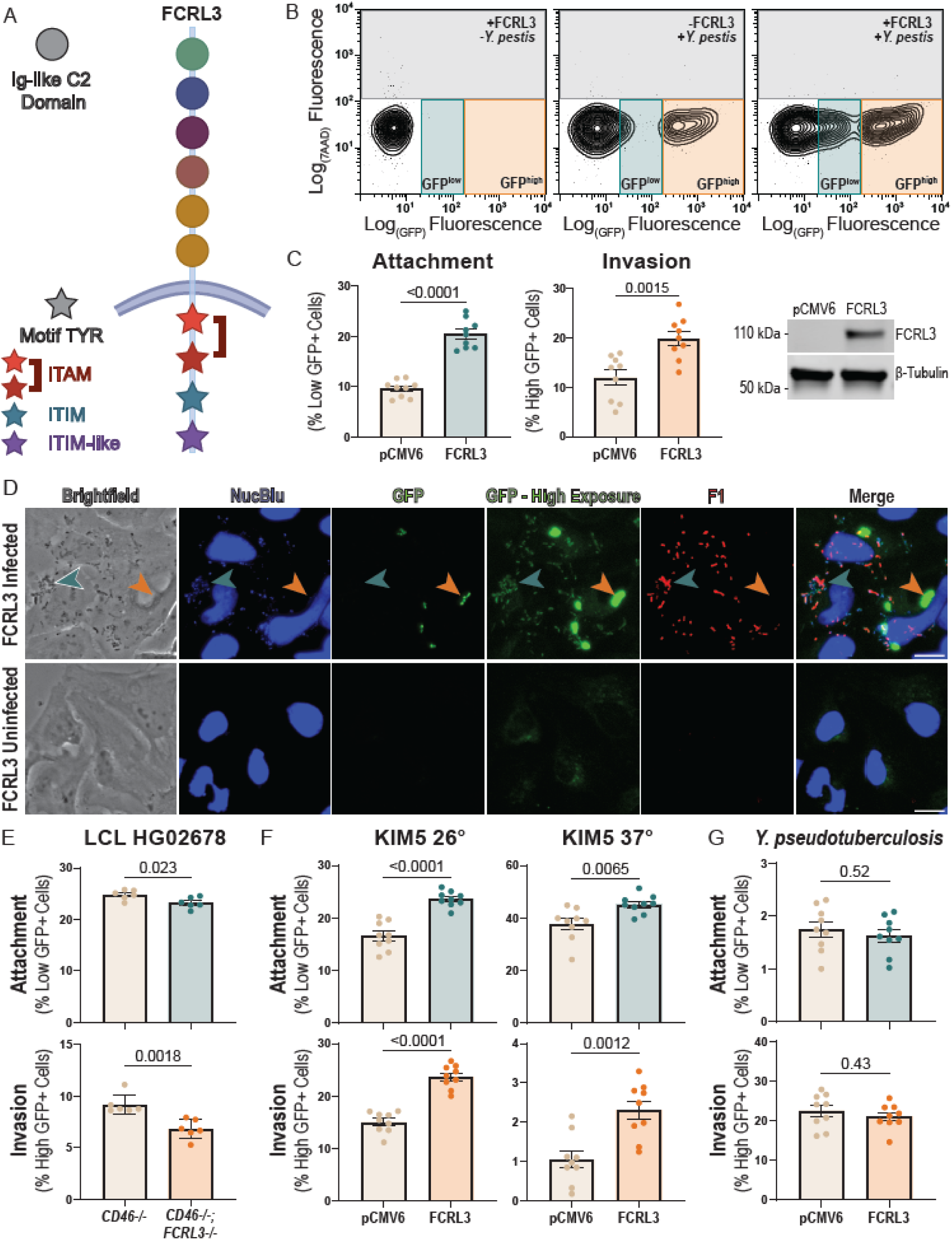
FCRL3 stimulates *Y. pestis* invasion. (**A**) Diagram of FCRL3 protein. Ig-like C2 domains are depicted as circles with the color indicating their previously established phylogenetic relationship, while tyrosines involved in intracellular signaling motifs are shown as stars. Red indicates an ITAM, light blue an ITIM, and purple indicates the ITIM-like/HemITAM Y722 motif in FCRL3. (**B**) Flow cytometry of *Y. pestis* infection of HeLa cells. HeLa cells (transfected with empty vector or FCRL3 plasmid) were infected for 1 hr with KIM6+ +pMMB67GFP, treated with gentamicin to kill extracellular bacteria for 1hr, and then GFP was induced in living, intracellular bacteria for 2 hrs with IPTG. Two populations of GFP+ cells were detected, consisting of HeLa cells infected with living, intracellular bacteria (GFP^high^, orange) in both conditions and those with attached, dead bacteria (GFP^low^, light blue) found in the *FCRL3* overexpression condition. (**C**) Overexpression of FCRL3 increases attachment (GFP^low^, light blue) and invasion (GFP^high^, orange). HeLa cells were transfected with empty vector or FCRL3 plasmid and assayed for attachment and invasion by flow cytometry as described in B. Three biological replicates in each of three experiments were plotted and an unpaired t-test was performed to determine significance. A western blot was performed to confirm the presence of FCRL3 after overexpression but not with empty vector using 1:200 FcRH3 antibody (Santa Cruz, C-2). (**D**) GFP^low^ cells have attached, extracellular *Y. pestis.* FCRL3 transfected HeLa cells were infected as described in B and fixed with 4% paraformaldehyde (PFA) after 4 hrs. Infected and uninfected cells were blocked with normal goat serum (but not permeabilized) and then incubated with 1:20 polyclonal anti-*Yersinia pestis* F1-Antigen antibody (BEI Resources, NR-31024) to stain extracellular bacteria. DNA was then stained with 1 drop/10mL of a membrane permeable dye NucBlue for nuclei visualization. To visualize the GFP^low^ bacteria, a high and low exposure GFP panel is shown. GFP^low^ bacteria are indicated with a teal arrowhead and GFP^high^ bacteria with an orange arrowhead. Cells were imaged using a 40× air objective on an EVOS M5000 Microscope. A 20µm scale bar is located on the merged image. (**E**) CRISPR-mediated knockout (KO) of *FCRL3* in LCL HG02678 causes significant decrease in attachment and invasion of *Y. pestis*. Cells were electroporated with either only *CD46* (control) or *FCRL3* + *CD46* guides and CRISPR Cas9. Pooled KO cells (>70% *FCRL3^-/-^*) were sorted for *CD46^-/-^* cells. Cells were assayed for KIM6+ *Y. pestis* attachment and invasion by flow cytometric gentamicin protection assay at 4 hpi. Three experiments with two biological replicates of each condition were plotted and an unpaired t-test was performed to determine significance. (**F**) *FCRL3* overexpression increases attachment and invasion of *Y. pestis* KIM5. HeLa cells transfected with empty vector or FCRL3 plasmid were assayed for KIM5 *Y. pestis* attachment and invasion by flow cytometric gentamicin protection assay at 4 hpi, after subculturing bacteria for 2 hrs and 40 min at either 26 or 37 degrees C. Three biological replicates in each of three experiments were plotted and an unpaired t-test was performed to determine significance. (**G**) FCRL3 overexpression has no effect on *Y. pseudotuberculosis* attachment and invasion. HeLa cells were transfected with empty vector or FCRL3 plasmid and assayed for attachment and invasion by flow cytometric gentamicin protection assay at 4 hpi. Three biological replicates in each of three experiments were plotted and an unpaired t-test was performed to determine significance. (**C, E-G**) Experiments were normalized by grand mean.

Previous studies have commonly used heterologous expression in non-phagocytic cells as evidence for the ability of cell surface proteins to function as phagocytic receptors (Herre et al., 2004; Uribe-Querol and Rosales, 2020; Zhang et al., 2006). To assess whether FCRL3 is sufficient to mediate attachment and invasion of *Y. pestis,* we overexpressed FCRL3 in HeLa cells, which have no detectable endogenous FCRL3 expression (**Fig. S3A**). Careful examination of these cells by flow cytometry revealed two distinct GFP+ populations after washing away non-adhered bacteria, defined as GFP^high^ and GFP^low^ (**Fig. 2B**). Overexpression of FCRL3 substantially increased both GFP^high^ and GFP^low^ populations (**Fig. 2C**). By fluorescence microscopy, we discovered the GFP^high^ population are host cells infected with living intracellular bacteria that are protected from gentamicin killing and express high levels of GFP in response to IPTG. These bacteria are found within a LAMP1+ *Yersinia*-containing vacuole (YCV) (**Fig. S3B**). We speculated that the GFP^low^ population were host cells with adherent, extracellular bacteria with low but detectable baseline GFP expression that had been killed after gentamicin addition. To confirm that these adherent GFP^low^ bacteria were indeed extracellular *Y. pestis,* we stained non-permeabilized cells using an antibody for the F1 capsule (found on the surface of *Y. pestis* after induction at 37°C (Du et al., 2002)). F1 antigen was detected exclusively on the surface of bacteria with low GFP expression (**Fig. 2D**). Thus, the GFP^low^ and GFP^high^ gates are measures of *Y. pestis* attachment and invasion, respectively, and overexpression of FCRL3 in HeLa cells caused significant increases in both phenotypes.

To determine whether FCRL3 is also necessary for attachment and invasion, we created a pooled knockout of *FCRL3* in LCL HG02678 using CRISPR-Cas9 (Bonglack et al., 2021; SoRelle et al., 2023). Disruption of *FCRL3* caused a moderate but significant decrease in attachment and invasion (**Fig. 2E**).

Following demonstration that FCRL3 is necessary and sufficient for invasion, we interrogated the conservation of FCRL3-mediated invasion across *Y. pestis* strains. KIM6+ is attenuated through loss of the pCD1 plasmid, so we tested whether FCRL3 could also stimulate attachment and invasion of KIM5, a strain that harbors pCD1 (Perry and Fetherston, 1997). KIM5 was grown under conditions that either stimulated (37°C) or suppressed (26°C) expression of the T3SS from pCD1. As expected, induction of the T3SS resulted in much lower levels of attachment and invasion, but FCRL3 overexpression still led to an increase in both phenotypes (**Fig. 2F)**. Thus, FCRL3-mediated invasion is independent of pCD1.

*Y. pestis* evolved from *Y. pseudotuberculosis,* an enteric pathogen, (Achtman et al., 1999), but has undergone both substantial gene loss and acquisition, including two *Y. pestis*-specific plasmids pMT1 and pPCP1 (Brubaker, 1991; Califf et al., 2015; Chain et al., 2004). To determine if FCRL3-mediated invasion was specific for *Y. pestis*, we infected HeLa cells overexpressing FCRL3 and measured the differences in attachment and invasion by flow cytometry. FCRL3 overexpression had no effect on *Y. pseudotuberculosis* invasion (**Fig. 2G**), indicating the specificity of this phenotype to *Y. pestis*.

### FCRL3 colocalizes with *Y. pestis* at sites of bacterial attachment

Following ligand binding, Fc receptors cluster together before activation of intracellular signaling (Li and Yu, 2021). We hypothesized that *Y. pestis* binding to FCRL3 might trigger a similar process and tested this through localization studies of overexpressed FLAG-tagged FCRL3 in HeLa cells. FLAG-FCRL3 protein primarily exhibited diffuse localization in uninfected cells. Remarkably, following exposure to *Y. pestis,* FLAG-FCRL3 colocalized with GFP^low^ (but not GFP^high^) *Y. pestis* (**Fig. 3**). This indicates FCRL3 undergoes clustering to sites of *Y. pestis* attachment, but that it is generally not present in the *Yersinia*-containing vacuole. Thus, we have demonstrated FCRL3-mediated attachment and invasion in HeLa cells with clustering of FCRL3 at sites of attachment, facilitating testing of mutant FCRL3 proteins and other members of the FCRL family.

**Figure 3.**
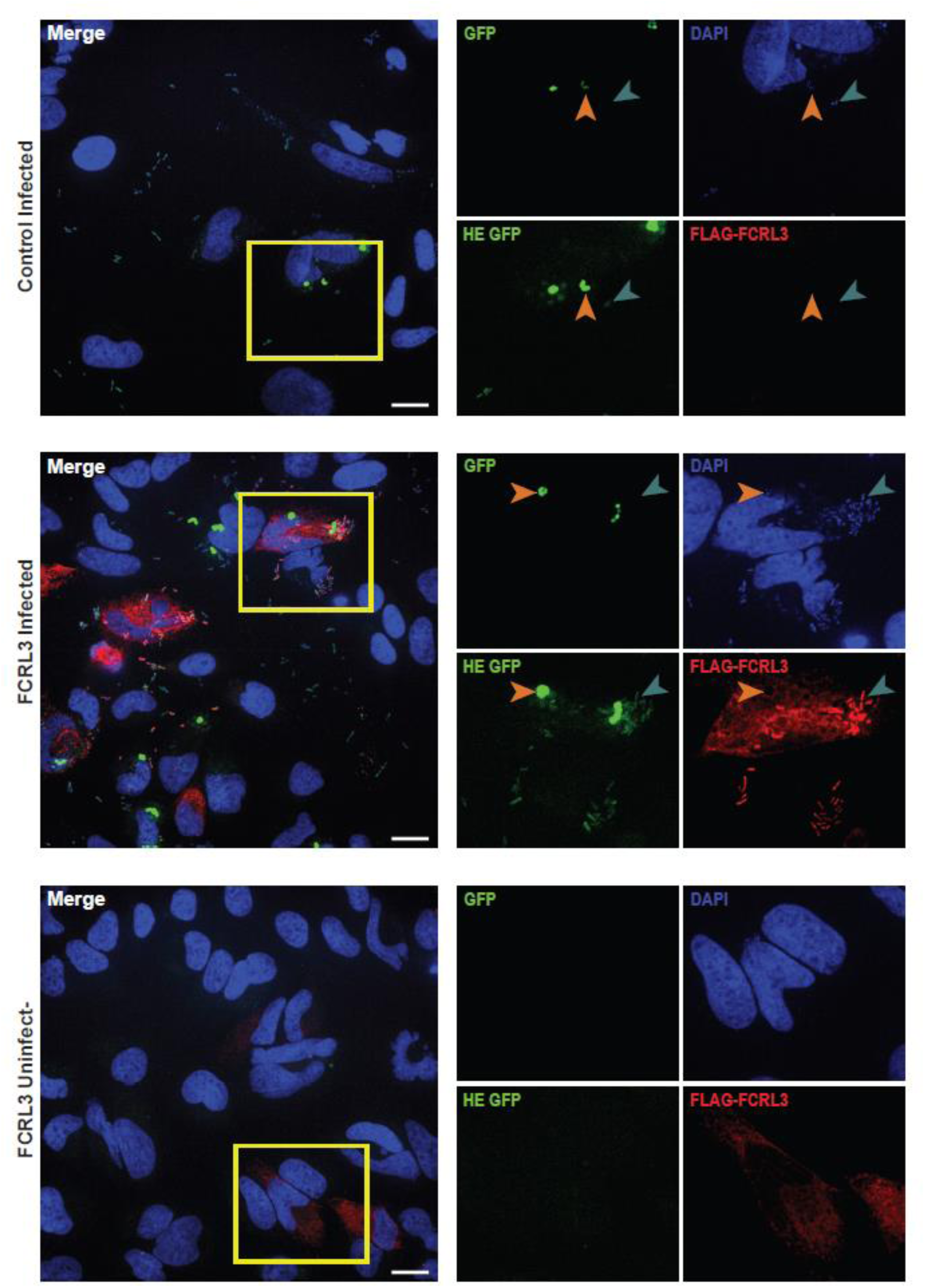
FCRL3 colocalizes with *Y. pestis* at sites of bacterial attachment. HeLa cells transfected with empty vector or FCRL3 plasmid were infected with KIM6+ +pMMB67GFP *Y. pestis* for 1 hr, treated with gentamicin for 1 hr, and induced with IPTG for 2 hrs prior to fixation with 4% PFA. After incubating for 30 min in block/perm, DNA was stained with 2.5µM DAPI, FLAG-FCRL3 was stained red using DYKDDDDK Tag antibody (Cell Signaling, D6W5B), and GFP+ bacteria are shown at low and high exposure to demonstrate the GFP^low^ bacteria. GFP^low^ bacteria are indicated with a teal arrowhead and GFP^high^ bacteria with an orange arrow head. Images were taken on a Zeiss Observer Z1 inverted microscope with a 63x water objective. A 20µm scale bar is located on the merged image.

### Functional redundancy for *Y. pestis* attachment and invasion among FCRL proteins

While FCRL3 overexpression in HeLa cells was sufficient to induce attachment and invasion, attachment or invasion were only moderately decreased in *FCRL3^-/-^* LCLs (see Figure 2E). We hypothesized that the presence of five other transmembrane FCRL family proteins in humans and hundreds of proteins in the immunoglobulin superfamily might lead to redundancy.

Each FCRL protein is structurally similar, containing a mix of five phylogenetically related types of extracellular Ig-like C2 domains, and intracellular motifs that mediate cell signaling (**Fig. 4A**, (Davis, 2007) (Li et al., 2014)). These proteins have considerable plasticity in the number, type, and location of these domains and motifs. We therefore hypothesized that this natural genetic variation might cause phenotypic variation in *Y. pestis* attachment and invasion. After overexpressing each available transmembrane construct (FCRL1, FCRL3-6)(Wilson et al., 2012) in HeLa cells and infecting, FCRL1 was the only FCRL protein unable to significantly increase attachment compared to an unrelated Ig-like domain containing receptor CD31 (PECAM-1) (**Fig. 4B**). The first and/or second Ig-like C2 domains of FCRL3 were present in all FCRL proteins that increased *Y. pestis* attachment (see Fig. 4A). Of those FCRLs that promoted attachment, only FCRL3 and FCRL5 significantly increased invasion (**Fig. 4B**). This property coincides with the presence of an ITAM (see Fig. 4A), which is reported to interact with the kinases SYK and Zap70 to activate cellular processes including endocytosis and phagocytosis (Xu et al., 2002). Interestingly, we found that while SYK overexpression alone had no effect on invasion into HeLa cells, co-expression with FCRL3 resulted in a significant increase compared to FCRL3 alone (**Fig. 4C**). Additionally, a SYK inhibitor (BAY 61-3606) inhibited *Y. pestis* invasion into LCLs (HG02678 and GM19204) (**Fig. 4D**). Thus, there are shared features among FCRL proteins that may mediate attachment (Ig-like domains 1 and 2) and invasion (an ITAM motif that signals through SYK), ultimately resulting in functional redundancy within the FCRL family.

**Figure 4.**
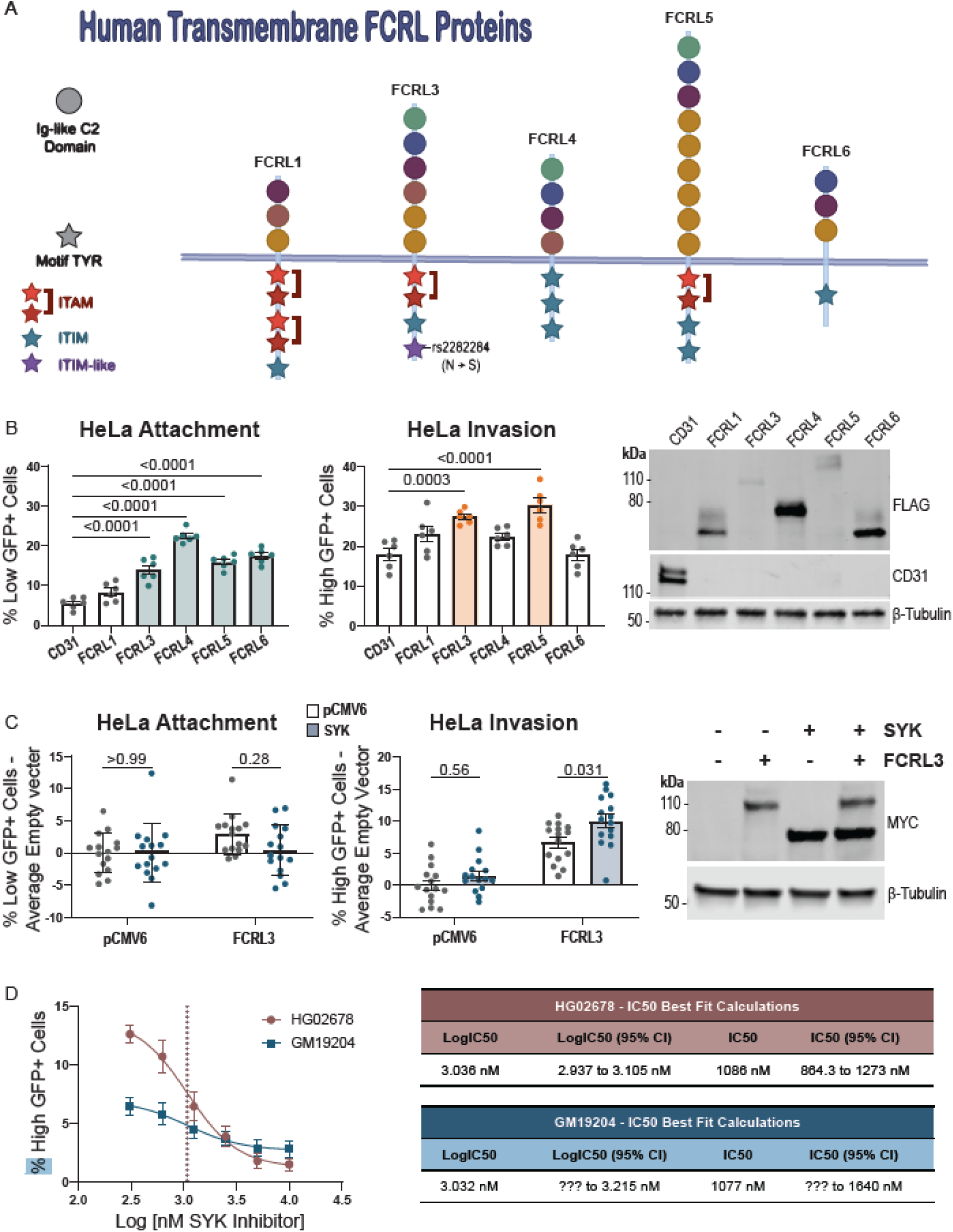
– Determinants of FCRL3 attachment and invasion reveal a critical role of FCRL3 domains and redundancy of function. (**A**) Protein homology of the human transmembrane FCRL proteins used in this study. Ig-like C2 domains are indicated by a circle and are colored by their phylogenetic relationship to one another. Tyrosine motifs are indicated by stars with color red, blue, or purple corresponding to ITAM, ITIM, or ITIM-like/HemITAM Y722 motif respectively. (**B**) Overexpression of other FCRLs have varying effect on attachment and invasion. HeLa cells were transfected with the indicated plasmid and assayed for attachment and invasion by flow cytometric gentamicin protection assay at 4 hpi. Colored boxes indicate FCRLs that are capable of increasing attachment (light blue) or invasion (orange). Two experiments with three biological replicates of each condition were plotted and a one-way ANOVA with Dunnett’s multiple comparisons test was performed to determine significance. A western blot was performed to show the presence of each construct using 1:1000 anti-FLAG (Sigma-Aldrich; M2) or 1:1000 anti-CD31 (Cell Signaling; 89c2). (**C**) Co-expression of SYK increases FCRL3-dependent invasion. HeLa cells were transfected with the indicated plasmids and assayed for attachment and invasion by flow cytometric gentamicin protection assay at 4 hpi. SYK co-expression is indicated with a blue shaded bar. Five experiments with three biological replicates of each condition were plotted and a two-way ANOVA was performed with Tukey’s multiple comparisons test to determine significance. Grand mean normalized values were used and the experimental average empty vector (pCMV6 only) value was subtracted from each value from each corresponding experimental replicate. Western blot displays the presence of MYC-SYK and MYC-FCRL3 by 1:200 anti-MYC-Tag (Cell Signaling; 71D10). (**D**) SYK inhibition significantly reduces invasion into LCLs. SYK inhibitor was added at 0.31, 0.63, 1.25, 2.5, 5, and 10 µM to LCL HG02678 and GM19204 at 60 min prior to infection with KIM6+. Cells were assayed for attachment and invasion by flow cytometric gentamicin protection assay at 4 hpi. The IC50 was determined for each line by nonlinear fit after log transforming the data. For GM19204, the lower confidence interval is undefined. Data are from three experiments of three biological replicates for each cell line and condition.

### Mutational analysis of FCRL-mediated attachment and invasion of *Y. pestis*

The FCRL proteins that can mediate binding (FCRL3-6) have Ig-like C2 domains with similarity to Ig like Domain (IgLD)-1 and/or IgLD-2 of FCRL3. Therefore, to determine if these Ig-like domains were required for binding, we deleted each as well as IgLD-3 for comparison. We observed a significant decrease in attachment relative to WT in all three mutants, but the effect was largest with IgLD-1 deletion. Additionally, only IgLD-1 deletion significantly decreased invasion (**Fig. 5A**). To further investigate this phenomenon, we inserted IgLD-1 into FCRL1, which does not cause a significant increase of attachment or invasion when overexpressed (see Fig. 4B). Insertion of IgLD-1 into FCRL1 was sufficient to significantly increase attachment and invasion, although not to the level of FCRL3 (**Fig. 5B**). The partial increase could be due to the lower protein expression of the mutant.

**Figure 5.**
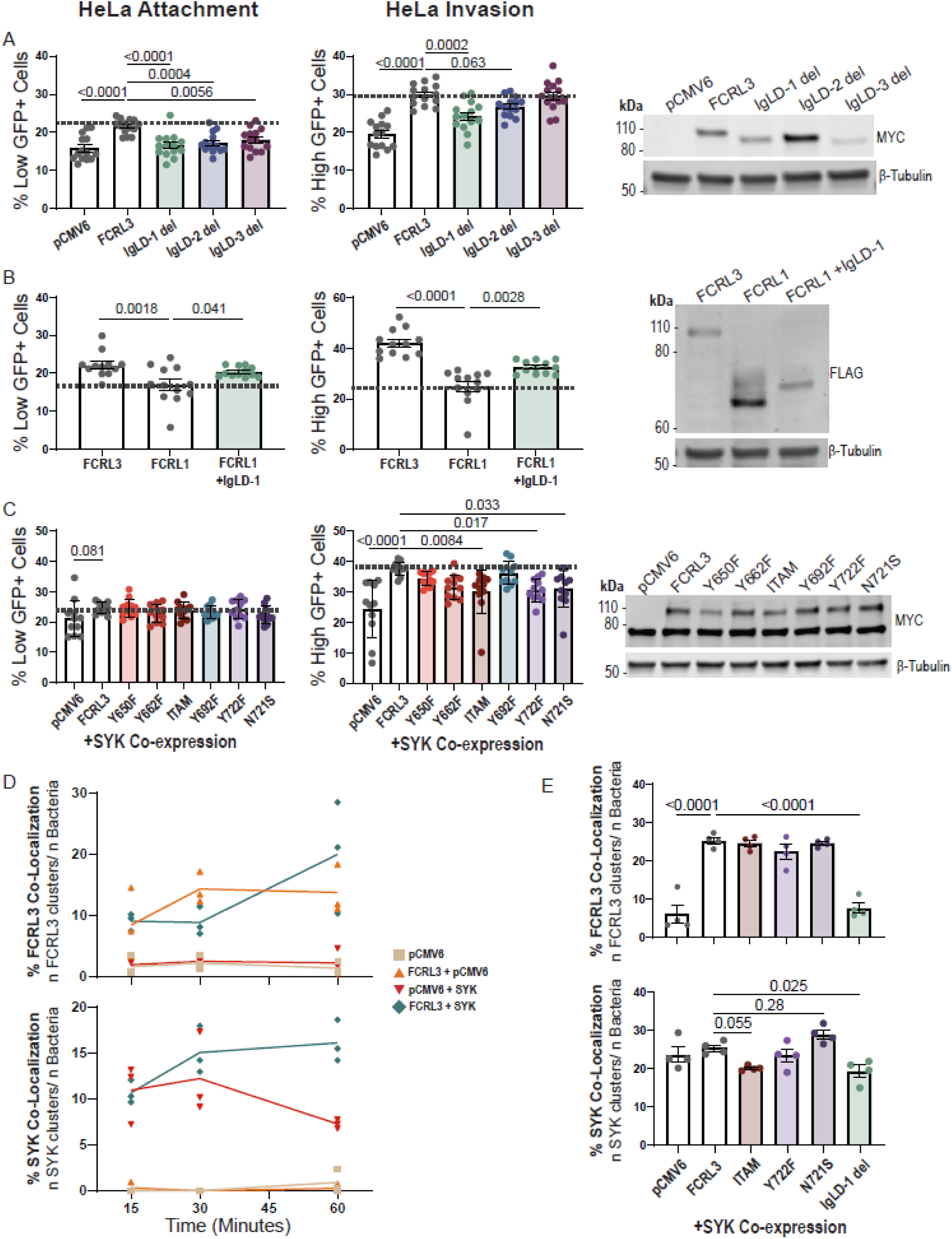
Mutational analysis of FCRL3 reveals the importance of Ig Like Domain 1 (IgLD-1) and the ITIM-like/HemITAM Y722 motif containing rs2282284 in attachment and invasion of *Y. pestis*. (**A**) Deletion of IgLD-1 leads to significant decreases in attachment and invasion. HeLa cells were transfected with the indicated plasmids and assayed for attachment and invasion by flow cytometric gentamicin protection assay at 4 hpi. Four experiments with three or five replicates were plotted and a western blot demonstrates each of the MYC-tagged constructs are overexpressed with 1:200 anti-MYC-Tag (Cell Signaling; 71D10). (**B**) Insertion of IgLD-1 into FCRL1 leads to an increase in attachment and invasion. HeLa cells were transfected with the indicated plasmids and assayed for attachment and invasion by flow cytometric gentamicin protection assay at 4 hpi. Four experiments with three replicates were plotted and western blot demonstrates each of the FLAG-tagged constructs are overexpressed using 1:1000 anti-FLAG (Sigma-Aldrich; M2). (**C**) Intracellular motif mutants lead to decrease in invasion when co-expressed with SYK. HeLa cells were co-transfected with the indicated plasmids and *SYK* plasmid then assayed for attachment and invasion by flow cytometric gentamicin protection assay at 4 hpi. Four experiments with two or three replicates for each condition are plotted. Western blot displays the presence of MYC-SYK (∼75 kDa) and MYC-FCRL3 (∼100 kDa) by 1:200 anti-MYC-Tag (Cell Signaling; 71D10). (**D**) Time course of FCRL3 and SYK recruitment to sites of GFP^low^ bacteria at 15, 30, and 60 min. HeLa cells transfected with the indicated plasmids were infected with KIM6+ +pMMB67GFP *Y. pestis* (pre-induced during liquid culture with IPTG 2 hrs prior to infection) and fixed with 4% PFA at the indicated time. Quantification of three experiments were plotted and ∼50 infected cells were counted for each condition. (**E**) Quantification of FCRL3 and SYK clusters at sites of GFP^low^ bacterial attachment for ITAM, Y722F, N721S, and IgLD-1 mutants. After infecting as described in D, cells were fixed at 1 hpi. Four experiments each quantifying ∼50 SYK+ cells from each condition were plotted. (**D-E**) Slides were stained with 1:400 anti-DYKDDDDK Tag (D6W5B) and 1:100 anti-SYK (4D10) (Cell Signaling). The experiment was performed blinded with one researcher transfecting the cells and another counting the cells after staining. (**A-C, E**) A one-way ANOVA was performed with Dunnett’s multiple comparisons test to determine significance. (**A-E**) Experiments were normalized using grand mean.

Downstream effects of Fc receptors are canonically mediated by phosphorylated ITAM or ITIM motifs and their interaction with kinases such as SYK (Getahun and Cambier, 2015). We found that the FCRLs with ITAM motifs were able to facilitate *Y. pestis* invasion (see Fig. 4B). Additionally, FCRL3 has a motif at Y722, that imperfectly matches the ITIM consensus (Kochi et al., 2009). FCRL3’s Y722 motif contains the missense mutation caused by rs2282284 (N721S), which is directly adjacent to the tyrosine in this possible HemITAM motif. To broadly test the importance of each tyrosine and motif to invasion, we created mutants containing tyrosine to phenylalanine mutations and a mutant with the rs2282284 N721S mutation. In HeLa cells, co-overexpression of the mutants constructs and SYK followed by infection revealed that the mutated ITAM (Y650F; Y662F) significantly decreased FCRL3-dependent invasion, the mutated ITIM (Y692F) had no effect, and Y722F (within the ITIM-like/HemITAM motif) caused the same reduction in SYK-mediated invasion as the mutated ITAM construct (**Fig. 5C**). In fact, simply altering the rs2282284 T allele to the minor C allele (N721S; associated with lower invasion) caused a similar reduction in invasion as Y722F. This indicates the Y722 motif promotes SYK and FCRL3-dependent *Y. pestis* invasion and the N721 position is critical for function. This finding also further suggests the Y722 motif may be acting as a HemITAM in FCRL3.

To test the importance of these mutations on FCRL3 clustering and SYK recruitment, we first monitored colocalization of FCRL3, SYK, and *Y. pestis* in HeLa cells overexpressing FCRL3 at 15-, 30-, and 60-min post infection. SYK and FCRL3 colocalization with attached *Y. pestis* increased throughout this time course when co-expressed (**Fig. 5D**). Therefore, FCRL3 clusters at sites of *Y. pestis* attachment and SYK is recruited to these sites. We then assessed the effects of the intracellular signaling (ITAM, Y722F, and N721S) and extracellular attachment (IgLD-1) mutations at the 1-hour timepoint. As predicted, the IgLD1 deletion led to a complete ablation of FCRL3 clustering, and a reduction in SYK recruitment. Interestingly, N721S resulted in a *greater* but not statistically significant fraction of SYK colocalized with FCRL3 and *Y. pestis* (**Fig 5E**). This was unexpected given the reduced attachment and invasion of this mutant, but we speculate that SYK can bind to N721S but is unable to efficiently trigger phagocytosis or other downstream steps.

### Direct binding of FCRL proteins to *Y. pestis*

While the FCRL family impacts attachment and invasion of *Y. pestis* into cells, this could be due to direct binding or secondary to effects of FCRLs on other cell surface molecules. To test this, we measured direct binding using the purified extracellular domain of FCRL5 fused to COMP5AP-AviTag-9xHis (Wojtowicz et al., 2020). The FCRL5 fusion protein was expressed at substantially higher levels than FCRL3, and overexpression of FCRL5 in HeLa cells was sufficient for stimulating attachment and invasion (see Fig. 4B, C), so it was used for direct binding experiments. The 9xHis tag facilitated purification of this secreted protein, while the human placental alkaline phosphatase (5AP) allowed for enzymatic detection of protein. A single-step affinity purification with cobalt beads resulted in high purity and 13-fold increase in 5AP specific activity (**Fig. 6A-B**). Following purification, FCRL5 fusion or negative control protein (CD31-COMP5AP-AviTag-9xHis) was incubated with *Y. pestis* for 30 min at 4°C, washed with high salt (600mM NaCl), and *Y. pestis* and bound proteins were lysed and resolved by SDS-PAGE. Due to high levels of endogenous alkaline phosphatase from *Y. pestis,* binding was measured by western blot. We observed that 38.8% of FCRL5 fusion protein bound to *Y. pestis* (4.6x more than to CD31; p=0.014) (**Fig. 6C**). Thus, *Y. pestis* directly binds to FCRL proteins to mediate attachment and invasion.

**Figure 6.**
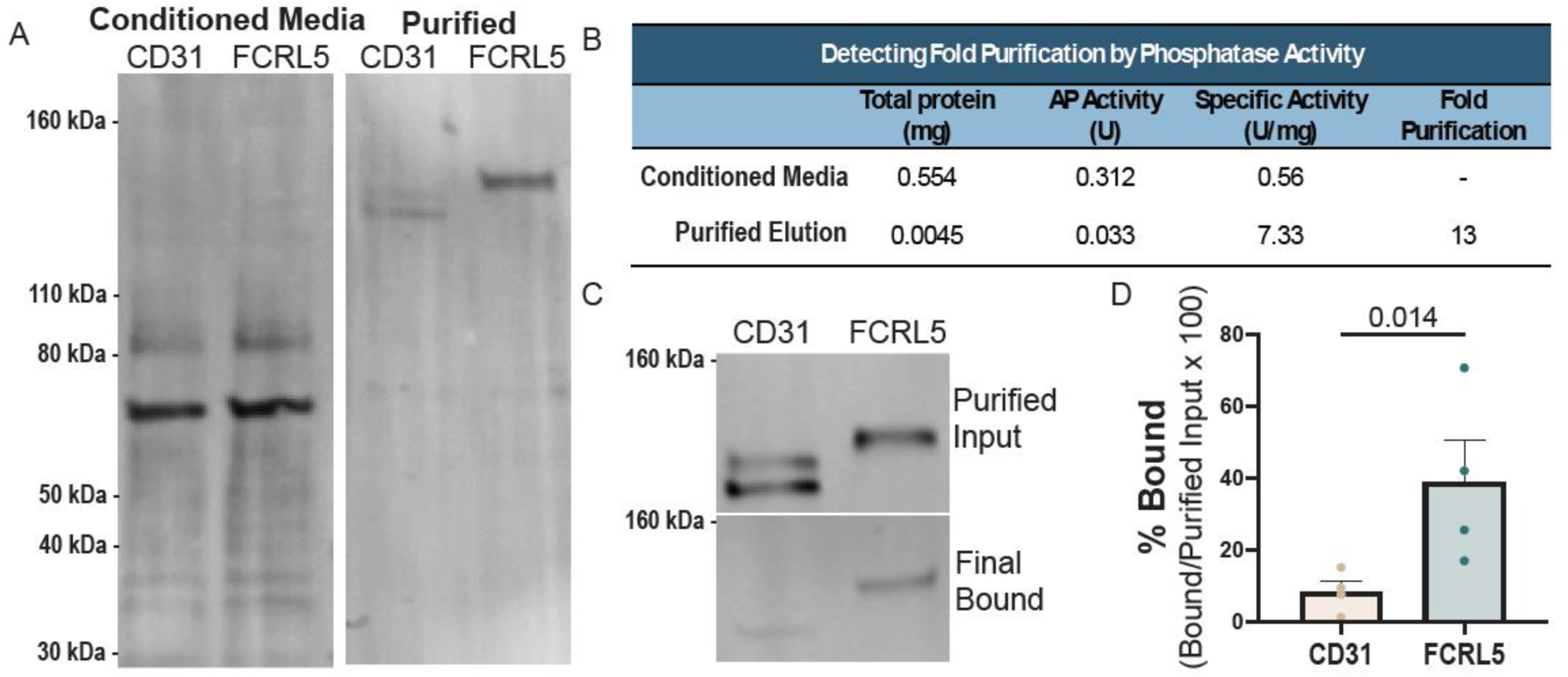
The FCRL5 extracellular domain directly binds to *Y. pestis*. (**A**) Total protein stain of supernatant from Expi293 cells transfected with CD31-COMP5AP-AviTag-9xHis or FCRL5-COMP5AP-AviTag-9xHis and purified protein following cobalt bead purification. (**B**) Table of purification demonstrates increased specific activity after purification. Total protein concentration was measured by Bio-Rad Protein Assay, and AP activity was measured using BluePhos® Microwell Substrate Kit, KPL. (**C**) Western blot of binding displays purified protein input and protein bound after incubation with *Y. pestis*, two washes, and subsequent lysis and sonication. Blots are stained with 1:1000 Avi Tag monoclonal Antibody (ThermoFisher; 1D11D10). (**D**) Bar graph showing quantification of bound vs. purified input western blots using infrared secondary antibodies (Li-COR Odyssey) from three binding experiments. Significance was tested by a paired ratio t-test.

### Impact of rs2282284 on other diseases: Chronic hepatitis C

While *Y. pestis* does not currently pose the threat to civilization that it has in the past, we hypothesized that rs2282284 might have additional consequences on current human health due to its role as an immune receptor. While there are no known genome-wide significant associations for rs2282284, PheWAS analysis of rs2282284 using BioBank Japan Pheweb (Nagai et al., 2017; Sakaue et al., 2021) showed a single trait with p-value less than expected by chance out of over 258 phenotypes: chronic hepatitis C virus infection (**Fig. 7A, B**; p=9.6×10^-5^; beta = –0.18; n=7110 cases, 169,588 controls) (Sakaue et al., 2021). The C allele (associated with reduced *Y. pestis* invasion) is associated with reduced risk of chronic hepatitis C. Colocalization analysis with COLOC (Giambartolomei et al., 2014) comparing the signals for *Y. pestis* invasion and chronic hepatitis C provided support for the two being due to the same causal variant (**Fig. 7C-E**; PP4=0.78).

**Figure 7.**
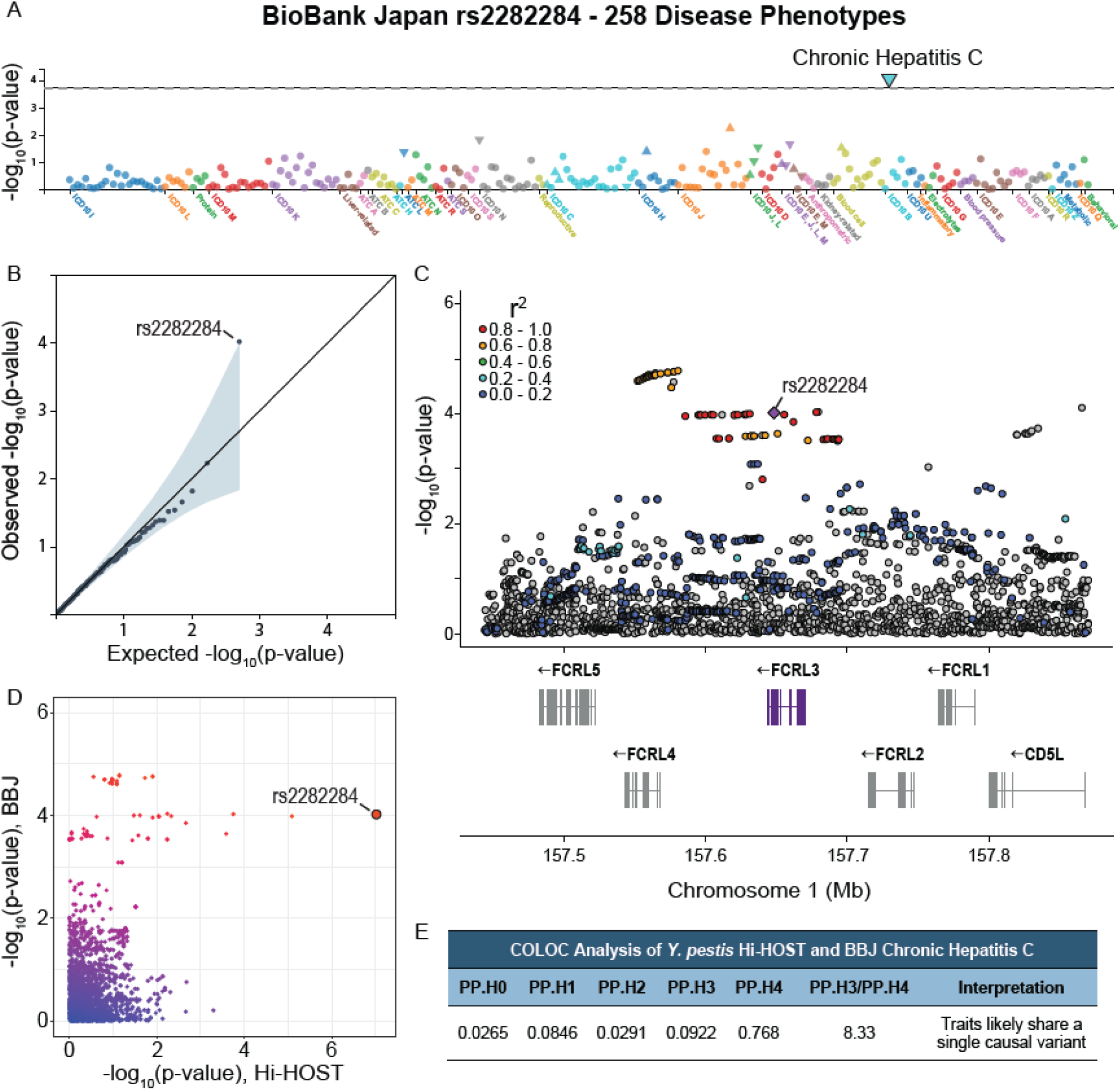
rs2282284 is associated with chronic hepatitis C. (**A**) Chronic Hepatitis C virus is associated with rs2282284. PheWAS plot displaying the –log_10_(p) values of 258 phenotypes in BioBank Japan PheWeb. Chronic Hepatitis C is indicated by a large blue triangle. Phenotypes are colored by disease group. (**B**) All 258 phenotype p-values were plotted on a quantile-quantile plot and rs2282284 was the only variant that significantly deviated from the neutral distribution (grey line). (**C**) Local Manhattan plot of the rs2282284 locus for the Chronic Hepatitis C virus phenotype. A purple diamond denotes rs2282284 and LD with SNPs in the locus is shown by red >= 0.8, orange = 0.6-0.8, green = 0.4-0.6, light blue = 0.2-0.4, dark blue < 0.2, and grey has no LD data. LD is based on JPT population LD. (**D**) Plot of the –log_10_(p-values) values from BioBank Japan (Y axis) and Hi-HOST (X axis) GWAS. rs2282284 is indicated by the large red dot. (**E**) COLOC analysis suggests both phenotypes have the same causal SNP. Table showing the posterior probabilities for each of 5 hypotheses with PP.H4 representing the posterior probability that both traits share the same causal variant. PP.H3/PP.H4 is > 5, making a stronger case for colocalization.

## DISCUSSION

Using a cellular GWAS of nearly a thousand LCLs from diverse populations and functional characterization, we have determined that FCRL proteins are direct binding receptors for *Y. pestis*. While FCRL proteins have recently been demonstrated to bind to immunoglobulins (Agarwal et al., 2020; Wilson et al., 2012), a direct role of these proteins in pathogen uptake has not been described. We report that FCRL-mediated uptake of *Y. pestis* consists of 1) binding of *Y. pestis* to the extracellular domain with a requirement for Ig-like domain 1; 2) clustering of FCRL at sites of attachment; and 3) SYK-dependent internalization mediated by ITAM and ITIM-like/HemITAM motifs. As these steps mirror how Fc receptors induce phagocytosis of opsonized bacteria (Uribe-Querol and Rosales, 2020), the question arises as to why *Y. pestis,* a highly adapted, human-specific pathogen would have developed this means of invasion after evolving from *Y. pseudotuberculosis* (Achtman et al., 1999), which we found does not utilize FCRL-mediated invasion. We speculate that during this evolution, *Y. pestis* has evolved mechanisms to hide from the immune system during lymphatic spread. While *Y. pestis* is phagocytosed by professional phagocytic cells, including after opsonization by Fc receptors (Ke et al., 2013), neutrophils and macrophages are intrinsically more bactericidal that lymphocytes. Therefore, direct FCRL-mediated phagocytosis of *Y. pestis* may provide an intracellular niche within B cells, some of the most abundant cells in the lymphatic system. Future studies will examine this further to determine whether this niche might facilitate dissemination or secretion of cytokines that alter infection with *Y. pestis*.

In addition to FCRL3, we found that other FCRL proteins also stimulate attachment and invasion of *Y. pestis*. Specifically, the presence of FCRL5, a paralog to FCRL3 with attachment and invasion phenotypes, may buffer the effect of the rs2282284 C allele in *FCRL3* on overall *Y. pestis* invasion—despite the mutation severely decreasing SYK-dependent FCRL3 function. This buffering effect of paralogs likely underlies the observation that human genes that have a paralog are less likely to have severe consequences in human disease (Hsiao and Vitkup 2008). The redundancy likely extends beyond the FCRL family, as the Ig-like C2 domain subtype required for FCRL3-mediated uptake (D1 from the nomenclature in (Davis, 2007)) is found not only in FCRL5, but also in conventional FCR family members (Davis, 2007). It remains to be determined whether the ability to facilitate *Y. pestis* binding and entry extends to other Fc receptors and indeed to the more distantly related members of the immunoglobulin superfamily. Previous reports describe two other FcRs that can bind bacteria in the absence of opsonization. CD89 appears to act as an innate immune receptor that binds and triggers phagocytosis and killing bacteria in macrophages (de Tymowski et al., 2019), while CD16A binds to *E. coli* to activate an ITAM-inhibitory pathway that prevents phagocytosis and contributes to sepsis (Pinheiro da Silva et al., 2007).

The SNP rs2282284 is present in all populations of the 1000 Genomes Project (Genomes Project et al., 2015) with a global MAF of 6% (from 2% (Han Chinese in Beijing) to 11% (Indian Telegu in the UK). Thus, this allele (the derived C allele associated with reduced *Y. pestis* invasion that encodes for the N721S mutation) was present prior to the out-of-Africa expansion, but there does not appear to have been obvious population differentiation at this locus or other signals of positive selection (such as singleton density score (SDS) (Field et al., 2016); best SDS = 1.40 p = 0.16 in African Ancestry Individuals from TOPMED dataset (Taliun et al., 2021)). Being near 5% MAF may have resulted in exclusion of the SNP in some GWAS, and even when included low numbers of homozygous C individuals may limit power to detect significant associations. Despite this, we found rs2282884 was associated with another infectious disease, chronic hepatitis C virus infection, in Japanese individuals. The same C allele that confers resistance to *Y. pestis* infection also confers protection against chronic hepatitis C. While hepatitis C virus primarily infects hepatocytes, it has been demonstrated to infect B cells (Chen et al., 2017), leading to the suggestion that hepatitis C virus may use B cells as a protective niche during chronic infection (Desombere et al., 2021). Thus, rs2282284 is pleiotropic and the derived C allele appears to protect against both *Y. pestis* and chronic hepatitis C, potentially in similar ways.

Due to the low global frequency despite seemingly positive consequences for historic humans, we speculate whether there might be balancing selective pressures on rs2282284 that have prevented this allele from reaching higher frequencies across the globe. These counterbalancing selective pressures might involve the reported inhibitory activity of FCRL3, whereby it interacts with the B cell receptor to inhibit activation (Kochi et al., 2009). With such a wide array of functionality, it is perhaps surprising that rs2282284 is not associated with more diseases. It has been found, however, that a different SNP, not in LD (rs3761959; LDlink all population r^2^= 0.0474), is an eQTL (GTEX (Consortium, 2020)) and protein-QTL (Suhre et al., 2017; Sun et al., 2018) for FCRL3. For rs3761959, the derived C allele (global allele frequency 54%) is associated with reduced FCRL3 expression and reduced risk of Graves’ disease (P= 2.27×10^−12^, (Zhao et al., 2013)) and rheumatoid arthritis (1×10^-10^, Europeans, (Ha et al., 2021)) (1×10^-8^, Europeans, (Kim et al., 2015)) (p=5.2×10^-6^, Japanese, (Kochi et al., 2005)), while increasing risk of multiple sclerosis (p = 1.0×10^-8^, European, (International Multiple Sclerosis Genetics et al., 2011) (International Multiple Sclerosis Genetics et al., 2019)). While this SNP is associated with autoimmunity, we observe no association of rs3761959 with *Y. pestis* invasion in our Hi-HOST dataset (p= 0.89), and modest association with chronic hepatitis C in the Biobank Japan dataset (p=0.044). These associations of rs3761959 and rs2282284 with different human traits may point to the role of the Y722 motif in a specialized subset of FCRL3 function. The rs2282284 SNP may specifically impair one aspect of phagocytosis or signaling important in these infectious diseases, while the overall expression level may be more important for functions involved in the development of autoimmunity.

In conclusion, we have used a cellular genome-wide association study of *Y. pestis* to reveal the FCRL proteins as previously unrecognized phagocytic receptors, regulated by human genetic variation that have pleiotropic effects on human infectious and autoimmune disease. We anticipate that future studies using Hi-HOST and *Y. pestis,* and other ancient and emerging threats, will reveal additional mechanisms in host-pathogen interactions and lasting consequences of pandemic pathogens on the human genome (Pittman et al., 2016).

## Methods

### Cell culture

LCLs were purchased from Coriell and cultured for eight days in RPMI 1640 media supplemented with 10% heat-inactivated fetal bovine serum (FBS), 2 mM glutamine, 100 U/mL penicillin-G, and 100 mg/mL streptomycin prior to assays. Cells were passaged at 150,000 cells/ml in 20ml total volume for three days after an initial 1-day rest period. LCLs were counted with a Guava Easycyte Plus flow cytometer (Millipore). HeLa cells were obtained from Duke University Cell Culture Facility and cultured in DMEM supplemented with 10% fetal bovine serum, 100 U/mL penicillin-G, and 100 mg/mL streptomycin. After 75-90% confluency was reached, cells were washed with PBS and lifted with 0.05% trypsin EDTA before neutralizing with FBS. HeLas were passaged at a 1:5 dilution in 10ml total volume for three days.

### *Yersinia* cellular infection

*Y. pestis* (KIM6+ and KIM5) and *Y. pseudotuberculosis* (IP32935) were tagged with an inducible GFP plasmid [pMMB67GFP from (Pujol and Bliska, 2003)]. Bacteria was grown overnight at 26°C in Heart Infusion Broth and ampicillin (100 µg/ml) then subcultured with a 1:33 dilution and grown for 2 hr 40 min at 37°C. Prior to infection, LCLs were washed once with RPMI (no additives) and then plated out in RPMI + 0.03% BSA at 100,000 cells/100 µl in a 96-well non-tissue-culture-treated plate. Invasion was conducted for 1 hr at a multiplicity of infection (MOI) of 30 after centrifugation at 100 x g for 5 min, followed by addition of gentamicin (50 µg/ml) for 1hr, and then culture was split into two separate cultures of 60 µl of cells with 140 µl of media to dilute gentamicin (15 µg/ml) and allow for collection at two timepoints. IPTG (1.4 mM) was added to turn on GFP expression for 120 min prior to 4 hr timepoints. 150 µl of cells were stained with 7-AAD (7-aminoactinomycin D; Enzo Life Sciences) and green and red fluorescence of 7000 cells was measured on a Guava Easycyte Plus flow cytometer (Millipore).

In HeLa cells, the same protocol was used replacing RPMI and RPMI + 0.03% BSA with DMEM (no additives) and plating either 12,500/100 µl in a TC-treated 96-well plate or 30,000/200 µl in a chamber well slide. Additionally, the media was replaced with DMEM media + 10% FBS, 15 µg/ml gentamicin, and IPTG (1.4 mM) for 120 min following 1 hr with high gentamicin.

### Genome-Wide Association

For each of 961 LCLs, three independent technical replicates were performed, and the mean was taken as the final invasion (4 hr % GFP+ cells) phenotype (see Table S1). Repeatability was calculated under a linear mixed model using the R package “*rptR*”, with 1000 bootstraps (for standard error and CI) and 1000 permutations (for permutation-based p-value).

Genotype imputation and genome-wide association analysis were performed as described in our previously published work (Gibbs et al., 2023; Wang et al., 2018b). Briefly, Hi-HOST was performed on 434 LCLs from HapMap project (CEU, YRI, CHB, JPT) (Consortium, 2005) and 527 LCLs from 1000 Genomes Project (ESN, GWD, IBS, KHV) (Consortium, 2010). Genotypes for the HapMap LCLs were obtained from HapMap r28 while genotypes for the 1000 Genomes Project LCLs were extracted from the 1000 Genomes Project Phase 3 genotypes (v.20130520). Both the HapMap and 1000 Genomes genotypes were imputed against the 1000 Genomes Project Phase 3 reference (GRCh37/hg19). Imputed genotypes were combined based on shared SNPs. A MAF filter of < 0.01 was applied, resulting in a total of 15,213,612 filtered SNPs.

Using the above genotypes, GWAS was conducted in PLINK v1.9 (Chang et al., 2015) using the QFAM-parents approach with adaptive permutation and a maximum of 10^9^ permutations. This approach performs linear regression to test for association while employing permutation of within– and between-family components separately to control for population stratification (Purcell et al., 2005). The human genome reference assembly (GRCH37/hg19) was used for all analysis.

### Phenotype– and SNP-based heritability analysis

Two different methods were applied to estimate heritability, as described for other Hi-HOST traits (Wang et al., 2018a). Briefly, the parent-offspring (PO) regression method estimated additive heritability using only phenotypic values and family relationships. Here, the slope is used as an estimation of heritability and was calculated at 0.1883. Secondarily, a genotype-based heritability estimate was performed using the GCTA GREML method (Yang et al., 2011). Here, SNPs with MAF of < 0.05 were excluded to construct the genetic relationship matrix (GRM). Zaitlen’s GREML method, which enables estimating GRM using both related and unrelated individuals, was used because of the family trio design used in Hi-HOST. The SNP-based *h*^2^ was 0.1881.

### Descriptive statistics and visualization

Descriptive statistics were performed with GraphPad Prism 10 (GraphPad Software, US) and with R Studio (Team, 2020). QQ plots were plotted using an adapted version of “Code Sample: Generating QQ Plots in R” (University of Michigan Center for Statistical Genetics, Matthew Flickinger) and Manhattan plots were created using “*fastman*” (Paria, 2022). Regional Manhattan plot were made using “*locuszoomr*” (Pruim et al., 2010). The size of each study or number of replicates, along with the statistical tests performed can be found in Figure Legends. All numerical data are presented as the mean ± SEM (standard error of mean).

### Cas9-RNP based editing of LCLs

To assemble RNP complexes, Synthego sgRNAs stocks for CD46 (a control transmembrane protein found on all nucleated cells) and FCRL3 at 30 μM and Cas9 at 20 µM were prepared (**Table S2**). In PCR tubes, 30 pmol of sgRNA and 10 pmol of Cas9 per guide were mixed and resuspension Buffer R was added to reach 7 µl total per reaction. While the RNP complexes incubated for 10-15 min at room temperature, 500,000 LCLs/reaction were centrifuged at 200 x g for 5 min, washed with PBS, and then suspended in Resuspension Buffer R at a concentration of 500,000 cells/5 µl. 5 µL of the cell suspension was added to each 7 µl of RNP complex mix (either CD46+FCRL3 or just CD46). Using the 10 µl Neon Transfection Kit, each reaction was pulsed at 1350 mV for 30 ms. The cells were then transferred into 200 µl of antibiotic free RPMI media + 10% FBS to rest for 48 hrs. After cells reached >2 million, they were stained with 200 ng/1 million cells PE CD46 (TRA-2-10) flow antibody (BioLegend) and the CD46 negative population was separated by fluorescence-activated cell sorting. Using Synthego suggested primers (**Table S2**), regions of interest were sequenced to ensure a frameshift causing a protein KO occurred.

### HeLa overexpression transfection

Plasmids (**Table S3**) were transfected using Lipofectamine 3000 per manufacturer’s instructions. 12,500/100 µl or 30,000/200 µl HeLa cells were plated on a TC-treated 96-well dish or in a chambered coverslip respectively in DMEM + 10% FBS and 1% pen/strep. After 24 hrs, the media was replaced with DMEM + 10% FBS without antibiotics after washing. After 1hr, 100 ng DNA, 0.2 µl of P3000 reagent, and 0.2 µl of Lipofectamine 3000 reagent were mixed with 10µl optiMEM and added to each 100 µl well or 2× for 200 µl wells.

### HeLa microscopy

Cells were washed with PBS and treated with 4% PFA at 1 or 4 hrs post infection for 15 min. Each well was then washed three times with PBS. 150 µl of PBS with either 1% saponin and 5% normal donkey serum (block/permeabilization solution) or only 5% normal donkey serum (block only, for non-permeabilized cells) was filtered and applied to the cells for 30 min. Primary antibody was applied at the indicated dilutions overnight at 4°C: 9e10 MYC 1:8 (Developmental Studies Hybridoma Bank), NR-31024 Polyclonal Anti-*Yersinia pestis* F1-Antigen 1:20 (BEI Resources), H4A3-s LAMP1 1:50 (Developmental Studies Hybridoma Bank) in block or block/permeabilization solution. The cells were washed three times with PBS and Alexa-fluor conjugated secondary antibodes (ThermoFisher Scientific) was added for 1 hr at a concentration of 1:2000.

DAPI was added to the cells in a concentration of 2.5µM for 5 min. The cells were washed three times with PBS and all liquid and the plastic guard was removed. Flouromount-G (Invitrogen) was added to each sample and a cover slip was added. The coverslip was left to dry for 24 hrs before imaging at 63x on a Zeiss Observer Z1 inverted microscope spinning disk confocal microscope or a EVOS M5000 Microscope. Final images were made using ImageJ.

### Quantification of Microscopy clustering

Slides were prepared as described above at 1 hpi after IPTG induction. Images were taken at 40x on an EVOS M5000 Microscope and ∼50 infected cells were counted for each condition. The counter was blinded to the conditions until after completion of counting.

### Construction of FCRL3 mutants

Deletion of Ig-like domains from FCRL3 were made using gene blocks and conventional cloning methods. Briefly, WT pCMV6-FCRL3 plasmid (RC214467 from OriGene) and 3 gene blocks (IDT), each containing two of the first three Ig-like domains (Q96P31, nucleotides 61-294 for Ig-like domain 1, 295-546 for Ig-like domain 2, and 574-810 for Ig-like domain 3 were cleaved with AsiSI and XbaI. Restriction-enzyme-digested WT FCRL3 plasmid lacking the first three Ig-like domains was resolved by gel electrophoresis and subsequent purification (Qiagen Gel Purification Kit). The restriction-enzyme-digested gene blocks were also purified (Qiagen PCR Purification Kit). Each digested gene block was ligated into the digested FCRL3 plasmid. The mutant FCRL3 plasmids were then transformed into E. coli (*NEB® 5-alpha Competent E. coli*), and successful genetic manipulation was confirmed with sequencing. Similarly, a gene block containing the Ig-like domain 1 in FCRL3 (nucleotides 61-294) was inserted into FCRL1 by restriction using AflIII and HindIII and insertion of the gene block by ligation.

To create point mutants in pCMV6-FCRL3, Lightning QuikChange (Agilent) was utilized according to manufacturer’s instructions. Primers were created with Agilent’s The QuikChange® Primer Design Program (**Table S4**). Mutations were validated by Sanger sequencing.

### Purification of FCRL5 extracellular domain and Y. pestis binding assay

For the extracellular domain of FCRL5, pD649-HAsp-FCRL5-COMP5AP-AviTag-9xHis (Plasmid #157554 from Addgene) was used. CD31 was again used as the control (pD649-HAsp-CD31-COMP5AP-AviTag-9xHis, Plasmid #157481 from Addgene). These plasmids were transfected into Expi293F cells using the ExpiFectamine 293 transfection kit per manufacturer’s instructions. Cells were split to 2.5-3×10^6^ viable cells/mL. After 24 hrs, 20mls of cells at high density (>5×10^6^ viable cells/mL) were split to a final density of 3×10^6^ viable cells/mL. For each transfection condition, 20000 ng DNA in 1000 µl of Opti-MEM™ I Reduced Serum Medium were mixed with 54 µl of ExpiFectamine™ 293 Reagent in 1000 µl of Opti-MEM™ I and incubated at room temperature for 10-15 min, then added to the seeded cells. 18-22 hrs post transfection, 120 µl of ExpiFectamine™293 Transfection Enhancer 1 and 1.2ml of ExpiFectamine™ 293 Transfection Enhancer 2 were added to the transfected cells. Conditioned media was collected 4 days post transfection.

For purification of His-tagged proteins in conditioned media, 12 ml of conditioned media was added to 10mg of Dynabeads™ magnetic beads. After a 2 hr incubation on a roller at 4°C, beads were washed four times with binding/washing buffer (50 mM SodiumPhosphate, pH 8.0, 300 mM NaCl, 0.01% Tween™-20). Beads were then incubated for 5 min in 200 µL of elution buffer (300 mM Imidazole, 50 mM Sodium phosphate pH 8.0, 300 mM NaCl, 0.01% Tween™-20). Purified supernatant was collected after the beads were applied to the magnet for 2 min.

For detection of direct binding, 3×10^7^ *Y. pestis* were incubated with 0.3% BSA in 60μl PBS pH 7.4 (1X) for 15-20 min at 4℃. After which, ∼1.8 μg of purified FCRL5 or CD31 control in 30 μl were added for a 90 μl reaction volume. After 30 min at 4℃, *Y. pestis* was spun down (5000 x g for 5 min), washed 2 x 5 min with 100 mM Sodium Phosphate, pH 8.0, 600 mM NaCl, 0.02% Tween™-20, and then lysed by boiling in 30 μl PBS pH7.4 (1X) 1x SDS-PAGE loading buffer for 5 min followed by sonication for 2 x 10 sec (Qsonica sonicator Q55, 1/8” Probe 15-20% amplitude). Inputs and bound fractions were resolved on a 4-20% gradient mini PROTEIN gel (Bio-Rad) and quantified by western blot probed with Avi Tag (1D11D10) monoclonal antibody (Thermo A01738-40, 1:1000 dilution) with quantification of bands using the western analysis function in LiCor Odyssey Image Studio Ver 4.0 to calculate the percentage of purified protein bound. Similar amounts of purified FCRL5 and CD31 were used in binding reactions based on quantification of purified proteins by western blot probed with anti-AviTag. In four binding experiments using two protein preparations, the relative amount of CD31 vs. FCRL5 used was 1.2 (+/-0.7).

### Western blot

Cell lysates were harvested from 100,000 cells using 30µl of TBS + 1% Octyl β-D-glucopyranoside (Sigma) and mini-cOmplete protease inhibitor table (Sigma; 1/4^th^ tablet added to 1.5ml of lysis buffer). Tubes were rocked at 4°C for 30 min before being centrifuged at 10,000 g for 5 min. Supernatant was added to 6x SDS loading buffer + βME and then boiled for 10 min. Samples were resolved on a 4-20% gradient mini PROTEIN gel (Bio-Rad) for 20 min at 80V then 40 min at 120V. Using the T77 semidry system (Amersham Biosciences (VWR)), proteins were transferred to a PVDF membrane using 60 mAmps. Blot was rinsed in PBS for 10 min and then placed in LICOR Odyssey Blocking Buffer for 1hr at room temperature. Primary antibody was placed in blocking buffer + 0.2% Tween overnight at 4°C. E7 beta Tubulin antibody (Developmental Studies Hybridoma Bank) was added to western blots at a 1:100 dilution for 1 hr at room temperature to confirm equal loading of samples. The membrane was washed four times for 5 min each in PBS + 0.2% Tween™-20 and IRDye secondary antibody (IRDye® 800CW Goat anti-Rabbit, IRDye® 680CW Goat anti-mouse IgG) (LiCor) was added 1:20,000 for 1 hr. The membrane was again washed four times for 5 min and then washed one more time in PBS before being imaged on the Licor Odyssey imager with Image Studio Ver 4.0.

### Colocalization analysis of Hi-HOST *Y. pestis* phenotype and BioBank Japan Chronic Hepatitis C GWAS

The R package “*coloc*” which is based on Giambartolomei et al.’s colocalization analysis (Giambartolomei et al., 2014), was used to determine if GWAS signals are due to a shared causal SNP. This method calculates the posterior probabilities that two traits are not associated in the locus of interest (PP0), only one trait is associated in the locus (PP1 and PP2), both traits are associated at the locus but with different, independent causal variants (PP3), or both traits are associated with a single causal variant in the locus (PP4).

For the Hi-HOST and BioBank Japan GWAS summary statistics, we filtered SNPs within a 500 kilobase (kb) window centered on rs2282284. The “coloc.abf” function was executed using the default prior parameters (p1 = 1×10^-4^, p2 = 1×10^-4^, and p12 = 1×10^-5^). PP4 between 0.700 to 0.900 indicated that the traits likely share a single causal variant. The PP4/PP3 indicated the intensity of the colocalization signal with values >5.00 further supporting colocalization.

### Resource availability Lead contact

Further information, as well as plasmids and bacterial strains generated for this study, are available by request from the lead contact, Dennis C. Ko.

### Materials availability

Plasmids and bacterial strains, as listed in the key resources table, are available upon request.

### Data and Code Availability

Summary statistics for Hi-HOST genome-wide association of *Y. pestis* invasion will be available upon publication from Duke Research Data Repository.

## Acknowledgments

We thank the investigators and individuals from diverse populations genotyped as part of the 1000 Genomes Project who have made their LCLs available through the Coriell Institute. We thank members of the Ko lab for useful discussion. RMK, SS, TD, LW, and DCK were supported by NIH grant R01AI118903. RMK and TD were supported by TriCEM Graduate Student Fellowships. RMK was supported by a Duke Precision Genomics Center student pilot grant. ML and NR were supported by R01CA140337.

## Author Contributions

Conceptualization, R.M.K., S.I.M., T.J.W. and D.C.K.; formal analysis, R.M.K., S.S., T.D., L.W., and D.C.K. investigation, R.M.K., S.S., T.D., L.W., and D.C.K.; funding acquisition, R.M.K. and D.C.K.; supervision, R.M.K, N.M.R., M.A.L, S.I.M, and D.C.K.; resources, R.M.K, S.S., N.M.R, M.A.L. and D.C.K.; writing – original draft, R.M.K., S.S., T.D., L.W., and D.C.K.; writing – review & editing, all authors.

## Supplementary information

Document S1. Figures S1-S3 and Tables S2-S4.

Table S1. Y. pestis invasion measurement for 961 LCLs (3 replicates and mean)

