## SupplementalFiguresTables for "Human genetic variation reveals FCRL3 is a lymphocyte receptor for *Yersinia pestis*"

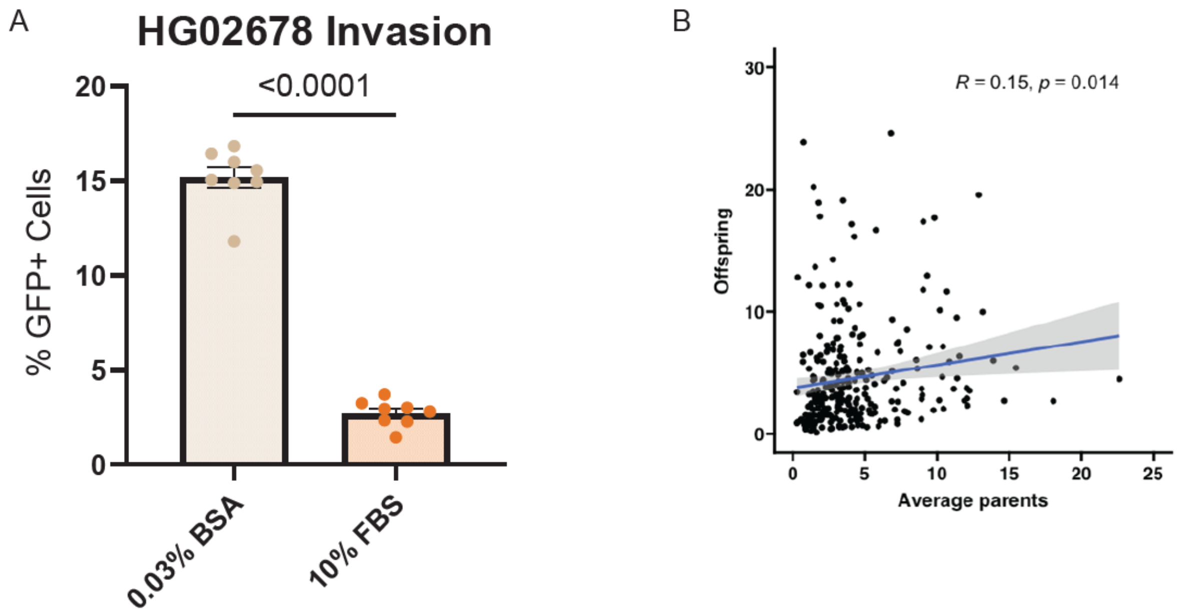


**Figure S1.** *Y. pestis* invasion into LCLs is inhibited by FBS and heritable. (**A**) Flow cytometric measurement of *Y. pestis* (KIM6+ +pMMB67GFP) invasion of LCLs at 4 hpi in RPMI media with either 10% FBS or 0.03% BSA after gentamicin protection assay. 2 experiments are grand mean normalized with 8 total replicates plotted for each condition. An unpaired t-test was performed to determine significance. (**B**) Parent-offspring linear regression of *Y. pestis* invasion into LCLs gives an estimated heritability of 0.15, a slope of 0.19 and a p-value = 0.014.

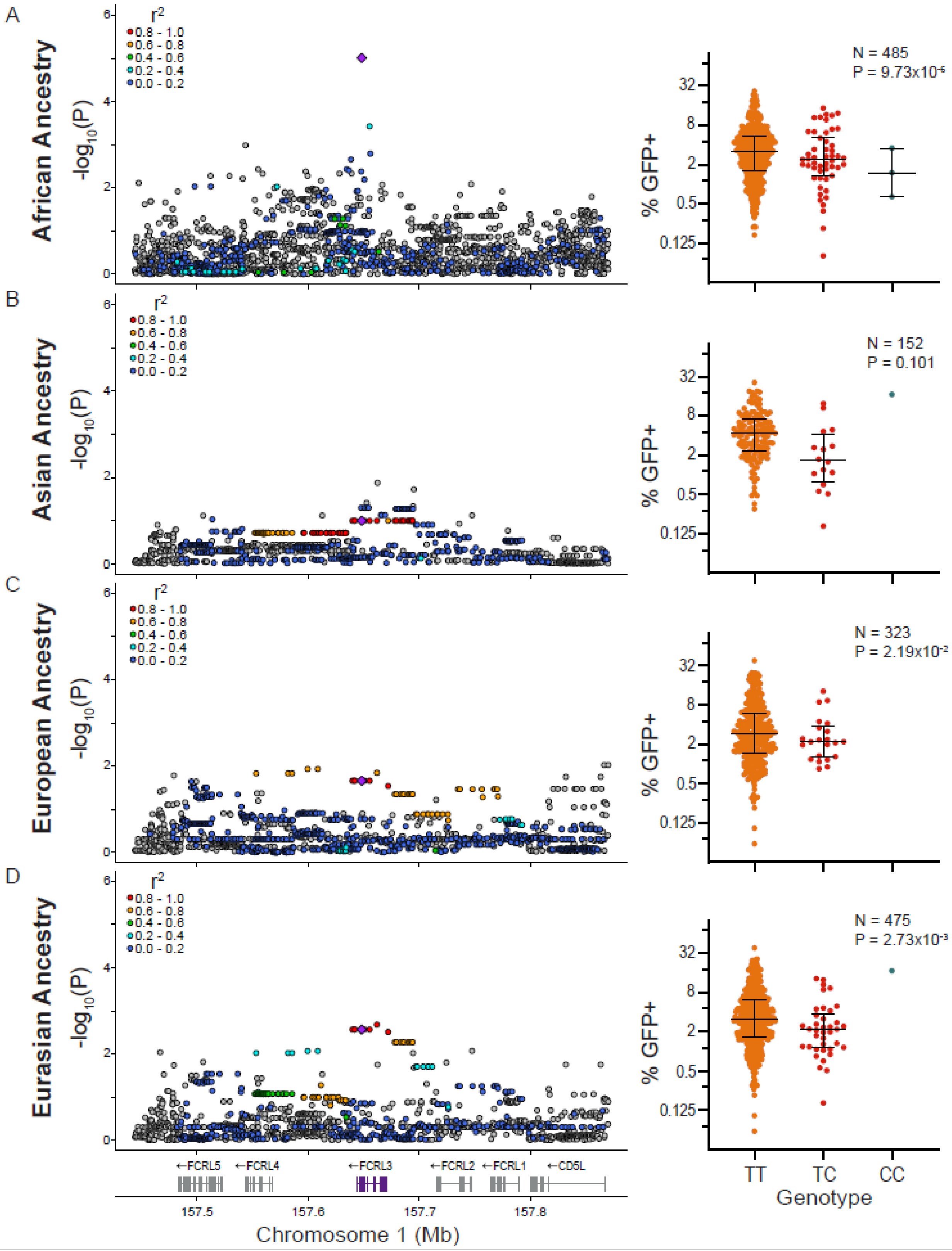

A
African Ancestry
-log10(P)
r2
0.8 - 1.0
0.6 - 0.8
0.4 - 0.6
0.2 - 0.4
0.0 - 0.2
N = 485
P = 9.73x10-6
% GFP+
B
Asian Ancestry
N = 152
P = 0.101
C
European Ancestry
N = 323
P = 2.19x10-2
D
Eurasian Ancestry
N = 475
P = 2.73x10-3
←FCRL5
←FCRL4
←FCRL3
←FCRL2
←FCRL1
←CD5L
157.5
157.6
157.7
157.8
Chromosome 1 (Mb)
TT
TC
CC
Genotype

**Figure S2**. Locus zoom and genotypic mean plots of LCLs stratified by ancestry. Local Manhattan plots and genotypic mean plots stratified by African (GWD, YRI, ESN), Asian (JPT, CHB, KHV), European (CEU, IBS), or EurAsian (JPT, CHB, KHV, CEU, IBS) ancestries. A purple diamond denotes rs2282284 and LD with SNPs in the locus is shown by red >= 0.8, orange = 0.6-0.8, green = 0.4-0.6, light blue = 0.2-0.4, dark blue < 0.2, and grey has no LD data. LD was determined by all populations within the stratified group. -$\log_{10}(p)$ values at the rs2282284 locus was plotted and phenotypes were plotted by genotype. All populations demonstrate the C allele of rs2282284 is associated with decreased invasion.

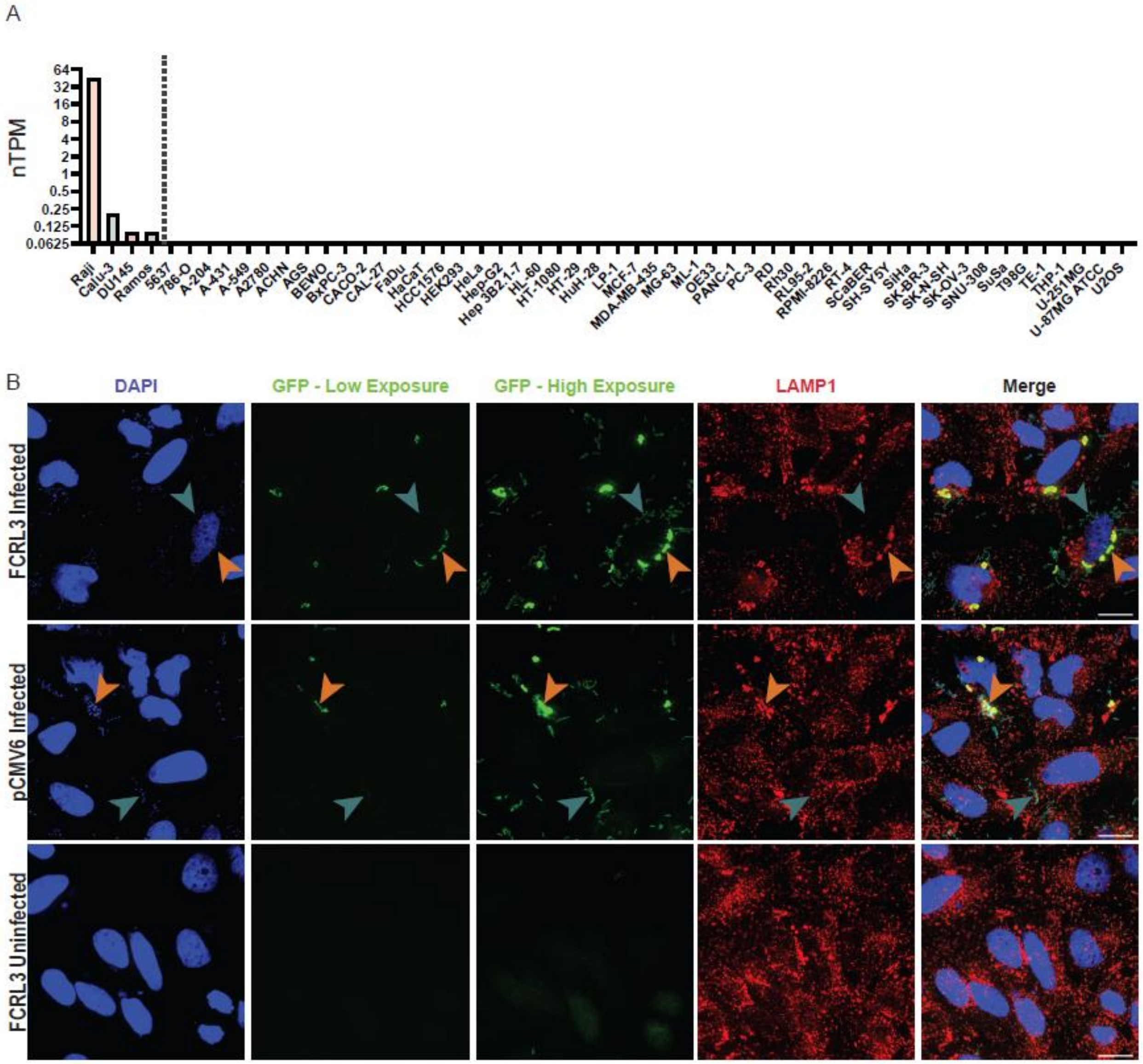


**Figure S3.** No FCRL3 protein is detected in HeLa cells and intracellular *Y. pestis* is in LAMP+ vesicles at 4 hpi. (**A**) Bar graphs of expression in common cell lines from the Human Protein Atlas (Uhlen et al., 2015) show expression only in lymphoma, Calu-3, and DU145 cell lines and no expression in HeLa. (**B**) GFP$^{high}$ cells have intracellular *Y. pestis* within LAMP1+ vacuoles. HeLa cells transfected with empty vector or *FCRL3* plasmid were infected with KIM6+ +pMMB67GFP *Y. pestis* for 1 hr, treated with gentamicin for 1 hr, and induced with IPTG for 2 hr prior to fixation with 4% paraformaldehyde. After incubating for 30 minutes in block/perm, DNA was stained with 2.5µM DAPI, LAMP1 was stained red with a 1:20 dilution of H4A3-s mouse LAMP1 antibody (Developmental Studies Hybridoma Bank), and bacteria are shown in green at high exposure to include the $GFP^{low}$ population. $GFP^{low}$ bacteria are indicated with a teal arrow head and $GFP^{high}$ bacteria with an orange arrow head. Images were taken on a Zeiss Observer Z1 inverted microscope with a 63x water objective. A 20µm scale bar is located on the merged image.

**Table S1.** (separate file) *Y. pestis* invasion measurement for 961 LCLs (3 replicates and mean)

**Table S2.** sgRNAs and primers for CRISPR mutagenesis and validation

| **GENE** | **GUIDE 1** | **GUIDE 2** | **GUIDE 3** | **FWD primer** | **REV primer** |
|---|---|---|---|---|---|
| **CD46** | GAGAAACAUGUCCAUAUAUA | AACUCGUAAGUCCCAUUUGC | UUGCUCCUUAGAGGAAAUAA | TGCCTGGGTGAATATGAATCTT | TGTCAGAAACAGCAAGTAGTTTTG |
| **FCRL3** | AAUUUCCAGGCUCUGUAAUU | GAUACCAAUAUGUGUCUCCC | CUGUGGACCAUGGAGGAUUG | TCTGCCTAGGATCCCTGCAT | ACCCTGGTCCTGACTGGA |

**Table S3.** Plasmids

| Plasmid backbone | insert | tag | Origin |
|---|---|---|---|
| pCMV6 | | MYC/FLAG | Simon Gregory |
| pCMV6 | FCRL3 | MYC/FLAG | Origene |
| pCMV-SPORT6 | CD31 | | Tim Wilson |
| pFLAG-CMV-4 | FCRL1 | FLAG | Tim Wilson |
| pFLAG-CMV-6 | FCRL3 | FLAG | Tim Wilson |
| pFLAG-CMV-7 | FCRL4 | FLAG | Tim Wilson |
| pFLAG-CMV-8 | FCRL5 | FLAG | Tim Wilson |
| pFLAG-CMV-9 | FCRL6 | FLAG | Tim Wilson |
| pD649-Hasp-COMP5AP | CD31 | AviTag/9xHis | Addgene |
| pD649-Hasp-COMP5AP | FCRL5 | AviTag/9xHis | Addgene |

**Table S4. Oligonucleotides for Quikchange mutagenesis**

| mutation | Forward | reverse |
|---|---|---|
| N721S | catgaagaagatgatgaagaaagctatgagaatgtaccacgtgta | tacacgtggtacattctcatagctttcttcatcatcttcttcatg |
| Y650F | ggagctggagccaatgttcagcaatgtaaatcctg | caggatttacattgctgaacattggctccagctcc |
| Y662F | ctggagatagcaacccgatttttcccagatctg | cagatctgggaaaaatcgggttgctatctccag |
| Y692F | gaggaacttacagtcctcttttcagaactgaagaagaca | tgtcttcttcagttctgaaaagaggactgtaagttcctc |
| Y722F | cccatgaagaagatgatgaagaaaactttgagaatgtaccac | gtggtacattctcaaagttttcttcatcatcttcttcatggg |
